# Disentangling intrinsic and extrinsic gene expression noise in growing cells

**DOI:** 10.1101/2020.08.26.268722

**Authors:** Jie Lin, Ariel Amir

## Abstract

Gene expression is a stochastic process. Despite the increase of protein numbers in growing cells, the protein concentrations are often found to be confined within small ranges throughout the cell cycle. Considering the time trajectory of protein concentration as a random walker in the concentration space, an effective restoring force (with a corresponding “spring constant”) must exist to prevent the divergence of concentration due to random fluctuations. In this work, we prove that the magnitude of the effective spring constant is directly related to the fraction of intrinsic noise in the total protein concentration noise. We show that one can infer the magnitude of intrinsic, extrinsic, and measurement noises of gene expression solely based on time-resolved data of protein concentration, without any a priori knowledge of the underlying gene expression dynamics. We apply this method to experimental data of single-cell bacterial gene expression. The results allow us to estimate the average protein number and the translation burst parameter.

## INTRODUCTION

Gene expression in all forms of life is subject to noise [1–7]. Experimentally, stochastic gene expression has been intensively studied, mostly in growing cells with exponentially growing cell volume [8–12] in which the copy numbers of mRNAs and proteins in general double on average during the cell cycle, as widely observed in bacterial and eukaryotic cells [8, 13–15]. To reduce cell cycle effects, a more biologically relevant protocol to quantify the stochastic degree of gene expression is to calculate the variability of *concentration* because most genes in proliferating cells exhibit approximately constant protein concentrations throughout the cell cycle over multiple generations [13, 16–21]. In yeast and mammalian cells, most genes also exhibit approximately constant mRNA concentrations throughout the cell cycle [14, 22, 23].

Considering the time trajectory of protein concentration as a one dimensional random walker in the space of concentration, it must be subject to an effective restoring force to prevent the divergence of concentration in the long time limit (note that cell growth contributes to this restoring force via the effect of dilution, as discussed extensively in Ref. [19]). However, little is known about how the strength of this restoring force is related to the stochastic nature of protein concentration. In this work we show that one can in fact infer the contribution of intrinsic and extrinsic noise (which we will define later) to the total gene expression noise from the properties of the restoring force. Previous works on solving this challenge often rely on particular models of the underlying dynamics of gene expression [24–27]. Here we develop a novel protocol which is, in contrast, insensitive to many of the details of the gene expression dynamics, and is thus applicable to a broad class of models. The protocol only relies on analysis of time-series data of protein concentrations. We expect it to be applicable to exponentially growing cells such as bacteria, yeast and cancer cells [8–12].

In the following, we first introduce a general framework to study the variability of mRNA and protein concentrations in growing cells. Within the framework, the initiation rates of transcription and translation can be age-dependent (here, we define age as the elapsed time since cell birth), *e.g.*, due to gene dosage effects as well as more complex cell cycle dependencies [15]. We show that independent of the details of the gene expression dynamics, the variances of mRNA and protein concentrations can always be decomposed into an extrinsic component and an intrinsic component. In the large cell volume limit, the intrinsic noise vanishes while the extrinsic noise remains finite [28]. We then introduce our protocol to extract the fraction of intrinsic noise, extrinsic noise and measurement noise in the total noise of protein concentrations and finally apply the method to experimental data of bacterial gene expression.

### Decomposition of noise

For simplicity, we consider a cell growing at a constant growth rate *µ* (Fig. 1). When the cell divides, the molecules are assumed to be segregated binomially and symmetrically between the two daughter cells [3]. Since for both bacterial and eukaryotic cells the degradation times of many proteins are longer than the cell cycle duration [29], we consider a non-degradable protein. Our results are equally valid for proteins with a finite degradation rate after some slight modifications (Supplementary Information, SI A). We allow the initiation rates of transcription and translation per cell volume, *k*_*1*_, *k*_*2*_, to be time dependent and, for example, they can exhibit stochastic dynamics. One can further express *k*_*2*_ = *β*_*m*_ where m is the mRNA concentration and *β* is the initiation rate of translation per mRNA. Mechanistically *β* is determined by the binding rate of ribosomes to mRNAs and largely determined by the concentration of ribosomes, which is roughly constant throughout the cell cycle [20].

**FIG. 1.**
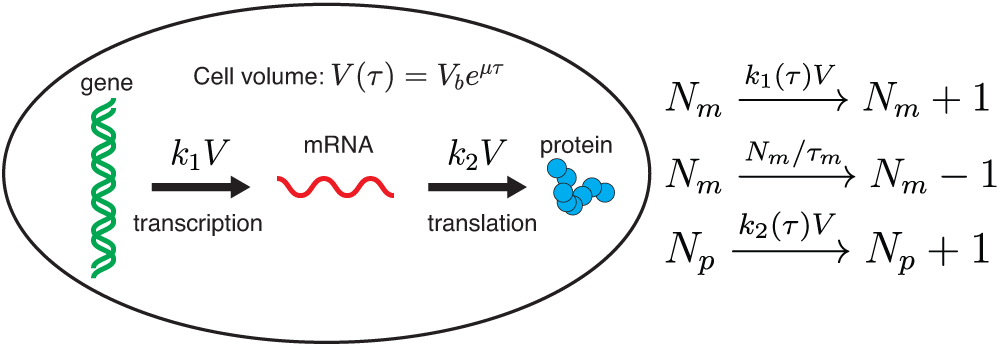
The cell volume *V* grows exponentially in time with a growth rate *µ* and *τ* is the cell age. *k*_1_ and *k*_2_ are the transcription rate and translation rate per cell volume which can be age-dependent. The chemical reactions of gene expression are summarized on the right. *N*_*m*_ and *N*_*p*_ are mRNA and protein numbers respectively. *τ*_*m*_ is the lifetime of mRNA using which one can define the translation burst parameter *βτ*_*m*_ (the average number of proteins produced in the lifetime of a single mRNA).

Consider an experiment where one tracks a single lineage of cells over multiple generations, records the data of protein concentrations *p* uniformly in time with resolution Δ*t*, and finally computes the resulting variance of concentrations based on all collected data. We find that the resulting variance of protein concentration 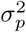 can be generally decomposed into three components (SI A):

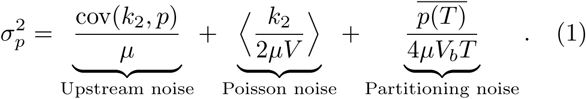

Here cov(*k*_2_; *p*) = 〈*k*_2_*p*〉 〈*P*〉 and 〈··〉 represents average over time. The first part represents the noise due to a fluctuating upstream factor, namely, the initiation rate of translation per cell volume. One important source of upstream noise is the fluctuation in mRNA copy number [28]. The second term represents the noise due to the stochastic production process which we denote as Poisson noise here. The last term stems from the random partitioning during cell division where *T* = ln 2/µ is the doubling time. The Poisson noise and the partitioning noise scale with the inverse of cell volume and their contributions to the variance 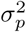 become negligible in the large cell volume limit conditioned on a fixed average concentration. In contrast, the upstream noise stems from the fluctuation in the translation rate per cell volume and it does not vanish in the large cell volume limit. We therefore define the sum of the Poisson noise and the partitioning noise as intrinsic and the upstream noise as extrinsic, consistent with previous works [28, 30]. We numerically confirm the validity of the noise decomposition for multiple gene expression dynamics including stochastic transcription and translation rate (SI B, Fig. S1).

We remark that the definition of extrinsic noise in our framework is different from the extrinsic noise inferred from the dual-reporter setup [1, 31], which is defined as the correlated noise of two identical genes controlled by the same promoters. The possible sources of extrinsic noise in the dual-reporter setup belong to a subset of those of the extrinsic noise in our framework which includes all possible upstream factors correlated or not across genes. Therefore, the extrinsic noise from the dual-reporter method is typically smaller than the extrinsic noise defined in our current framework, as we will discuss further later.

### Extracting the fraction of intrinsic and extrinsic noise

In the following, we discuss a protocol to disentangle the contribution of intrinsic and extrinsic noise to the total noise based on the time trajectory of concentration (Fig. 2a, b). We consider a discrete increment of protein concentration over a small time window, Δ*p*(*t*) = *p*(*t* + Δ*t*) − *p*(*t*), which can be expressed as

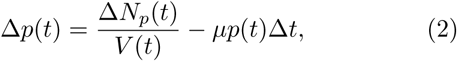

where Δ*N*_*p*_(*t*) is a random variable from a Poisson distribution with mean *k*_*2*_(*t*)*V* Δ*t* assumed constant within the small time interval Δ*t*. The second term on the right side arises from dilution due to cell growth. The protein concentration fluctuates but does not diverge in the long time limit, therefore we can make an analogy with a Brownian particle attracted to a fixed point with a linear restoring force equal to −*kx* where *k* is the spring constant and x is the particle position relative to its equilibrium point. In the case of a Brownian particle, one can find the spring constant of the restoring force as the slope in the linear fitting of the discrete velocity Δ*x*/Δ*t vs*. *x*. In the case of protein concentration, one can do a similar analysis by linearly fitting the protein production rate Δ*p*/Δ*t vs*. *p*. Considering a least square linear fitting, the slope of the linear fitting is found to be

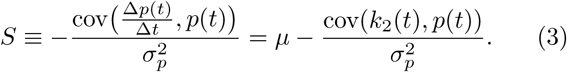

where we have used Eq. 2. If the covariance between the translation rate and protein concentration vanishes, the spring constant of the restoring force is simply the growth rate. Combined with Eq. 1, we find that the slope is proportional to the growth rate and the proportional constant is precisely the fraction of intrinsic noise in the total protein concentration noise variance:

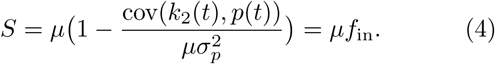

**FIG. 2.**
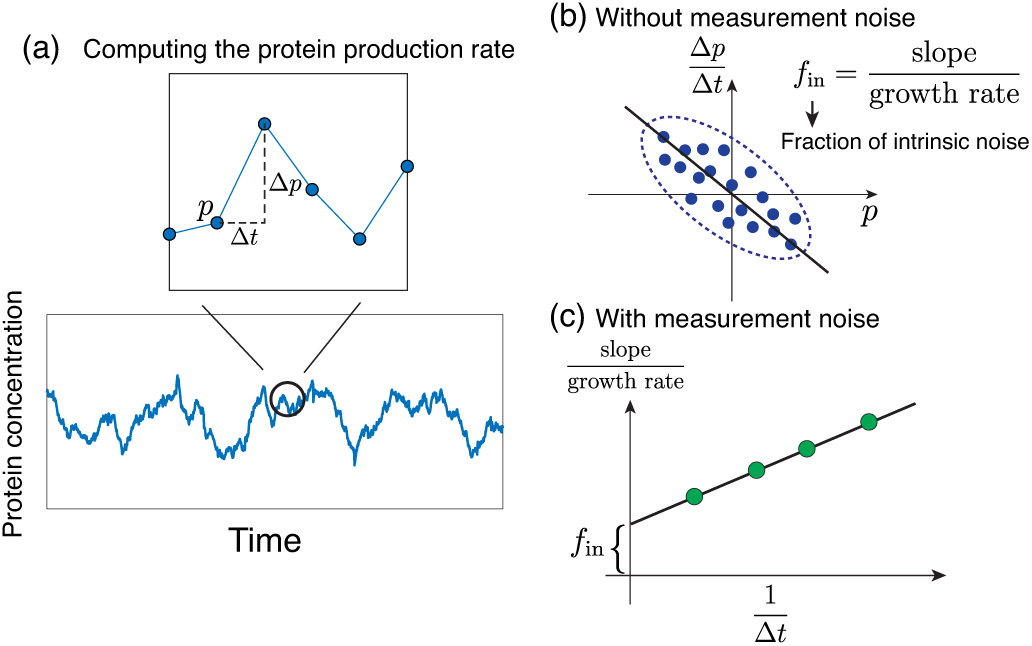
(a) Given a time series of protein concentration, we first compute the protein production rate as the discrete time derivative of protein concentration with a time interval Δ*t*. (b) Next, we perform a linear fit of the protein production rate Δ*p*/Δ*t* against the current protein concentration p and consider the absolute value of the fitted slope. In the case of negligible measurement noise, the fraction of intrinsic noise is the ratio between the slope and growth rate. (c) For experimental data with measurement noise, we compute the protein production rate for multiple time intervals Δ*t* and repeat the protocol in (b) for each time interval. Finally, we perform a linear fit of the normalized slopes against 1/Δ*t* and infer the fraction of intrinsic noise from the intercept.

The above equation shows that we can extract the fraction of intrinsic noise *f*_in_ in the total noise simply by linearly fitting the time derivative of the protein concentration against the current protein concentration without any a *priori* knowledge of the underlying gene expression dynamics. Extrinsic noise reduces the slope in the linear fitting which precisely equals the growth rate *µ* in the absence of extrinsic noise. An extended discussion along with an intuitive argument on the effects of extrinsic noise based on a Langevin equation is provided in SI F. We remark that our protocols are also applicable to nongrowing cells with a constant cell volume given the lifetime of the studied protein is known (SI A).

### Analysis of synthetic data

We test Eq. 4 on synthetic data, first considering a constitutively expressed gene where the initiation rate of transcription per cell volume *k*_1_ is constant as is the initiation rate of translation per mRNA *β*. This assumption corresponds to the case in which both RNA polymerase and ribosomes are limiting for gene expression, as discussed in detail in Ref. [19]. We compute *f*_in_ numerically using Eq. 1 and compare it with the prediction from Eq. 4, finding excellent agreement (Fig. 3a). To test the robustness of our protocol, we also verify our theoretical results on various other gene expression dynamics: (1) the scenario of transcriptional bursting where a gene switches from “off” state to “on” state with rate *k*_on_ and vice versa with rate *k*_off_ (Fig. 3b); (2) a gene with a constant transcription rate proportional to the gene number which doubles in the middle of the cell cycle (Fig. S2a); this scenario corresponds to the situation when the gene copy number is the sole limiting factor of transcription [19]; (3) a gene with a transcription rate modulated throughout the cell cycle due to a finite period of DNA replication (Fig. S2b, see details in SI E); (4) a gene with a fluctuating transcription rate (Fig. S2c); (5) a gene with a fluctuating translation rate per mRNA (Fig. S2d). In all cases, the predicted fractions of intrinsic noise match the actual values well. We also find that in all cases increasing the translation rate per mRNA *β* increases the fraction of extrinsic noise as the effects of upstream noise are amplified, consistent with the analytical results of constitutively expressed genes (SI C, D).

**FIG. 3.**
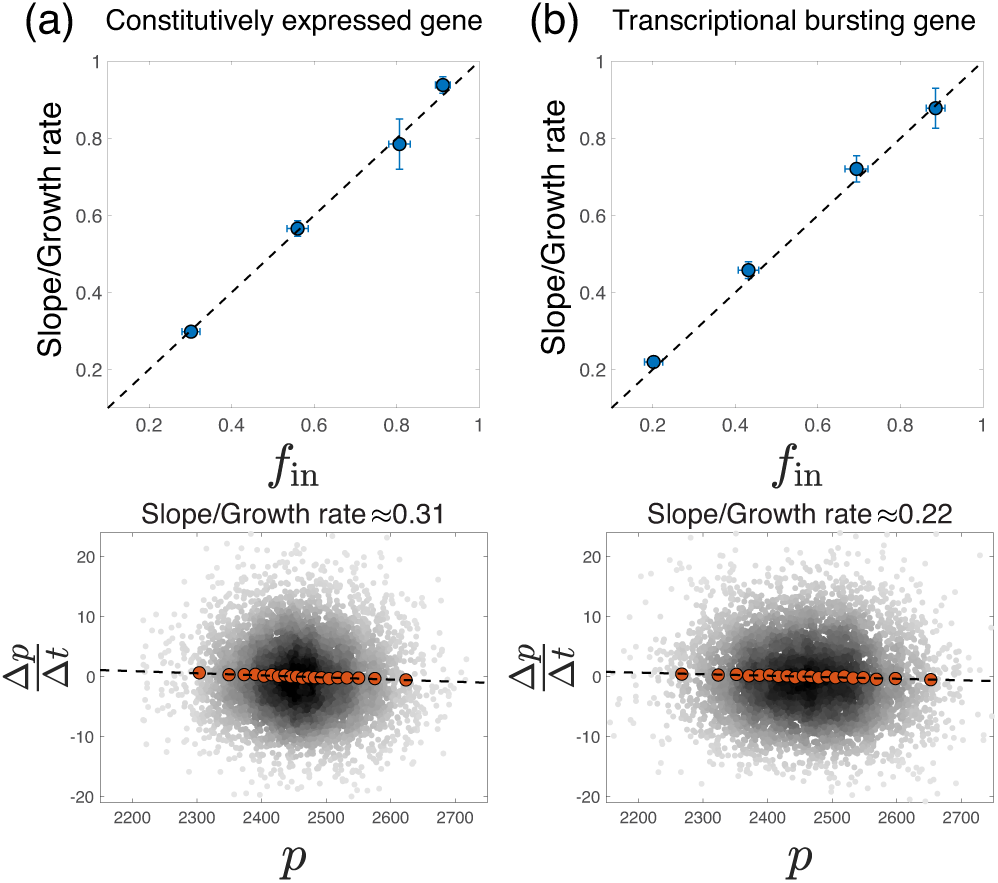
(a) Simulation of a constitutively expressed gene. (Upper) We compare the predicted fraction of intrinsic noise (*y* axis) to the measured value (*x* axis). (Bottom) An example of the raw numerical data with the binned data shown as well (red circles). The dashed line is the linear fit of the raw data. The same analysis also applies to panel (b). Here *k*_1_ = 10. (b) Simulation of a transcriptional bursting gene with *k*_on_ = 10, *k*_off_ = 10, *k*_1_ = 20. In all upper panels, the doubling time *T* = 60, *τ*_*m*_ = 10, and *β* is varied so that log_10_ *β* = −2;−1.5;−1;−0.5. In all bottom panels, log_10_ *β* = −0.5. We compute the time-derivative of protein concentration with a time interval Δ*t* = 0.5. The errorbars are computed as the standard deviation of 5 independent simulations.

In our framework the extrinsic noise is extracted from the time trajectory of the protein concentration of a single gene, which is distinct from that of the dual-reporter method. If the two genes in the dual-reporter setup share the same uctuating translation rate *k*_2_(*t*), the two definitions of extrinsic noise will coincide (SI G, Fig. S5a). However, if the correlated noise between the two genes is at the transcriptional level, the extrinsic noise inferred from the dual-reporter will be smaller than the one extracted from our protocol, which we confirm numerically (Fig. S5b).

### Analysis of experimental data

Experimentally, the measured protein concentration is always augmented by measurement noise. To model the effects of measurement noise, we assume the measured protein concentration at time *t* to equal

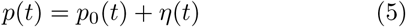

where *p*_0_(*t*) is the actual protein concentration and ƞ(*t*) is the measurement noise term assumed uncorrelated between different measurements. We will revisit this assumption later on and show that the datasets we analyzed are consistent with it. The covariance between Δ*p*/Δ*t* and *p* becomes 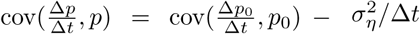. Compared with Eq. 4, the slope in the linear fitting of Δ*p*/Δ*t vs. p* is modified to

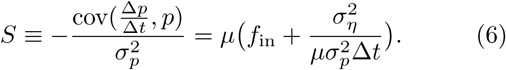

We confirm Eq. 6 using numerical simulations with artificial measurement noise. In this case since 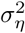 is assigned and *f*_in_ is known, we can directly compare the left and right sides of Eq. 6, obtaining good agreement (SI H, Fig. S6). Experimentally, the fluorescence level may not accurately reflect the instantaneous protein number due to a finite maturation time of the fluorescent protein. We have confirmed that the effects of a finite maturation time does not affect our results for experimentally relevant values of the maturation times [33] (SI I, Fig. S7).

We analyze two datasets of *E. coli* growth. In both, cells are exponentially growing and a fluorescent protein is constitutively expressed [8, 32]. A single lineage of cells is tracked for about 100 generations with cell volume and fluorescence level measured simultaneously. In both cases, the time interval between two consecutive data points is 1 min. To compute *f*_in_ for the experimental data, we increase the time interval to compute Δ*p*/Δ*t* and find the slopes in the linear fitting of Δ*p*/Δ*t vs. p* for each time interval (see examples for Δ*t* = 1 min in Fig. 4A, B). We then linearly fit the resulting slopes as a function of 1/Δ*t* (Fig. 2C) and the results agree well with the prediction of Eq. 6 (Fig. 4C). Notably, this allows us to infer both *f*_in_ as the intercept of the linear fit, and the fraction of measurement noise from the slope. The results are summarized in Fig. 4D. To justify the assumption of uncorrelated measurement noise, we show that the scaling with Δ*t* in Eq. 6 is violated for correlated measurement noise (SI H, Fig. S6).

**FIG. 4.**
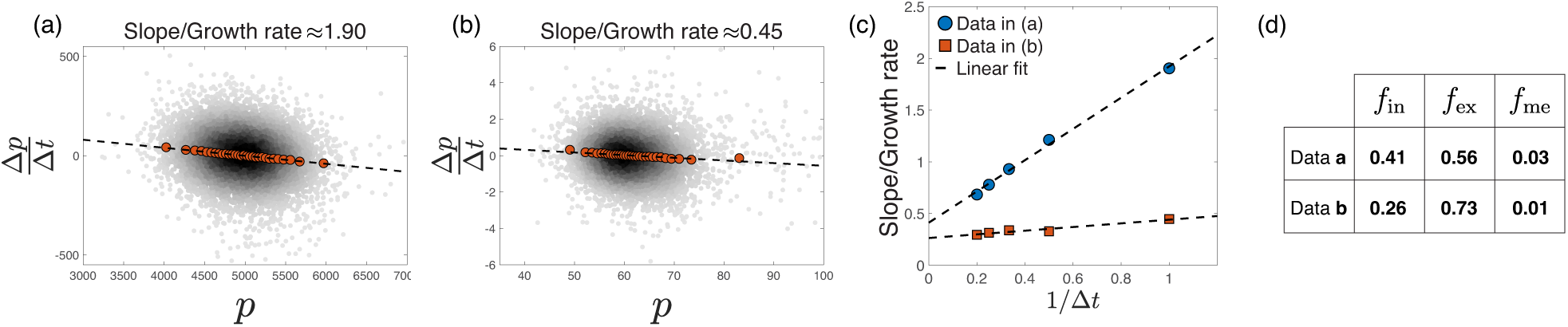
(a) We compute the time derivative of protein concentration as a function of the current protein concentration using data from Ref. [32] and the measured slope normalized by the growth rate is 1.90. The time interval used is Δ*t* = 1 min and the growth rate is *µ* = 0.0213 min^−1^. (b) We repeat the analysis using another data from Ref. [8] where the measured slope normalized by the growth rate is 0.45. Here Δ*t* = 1 min and µ = 0.0327 min^−1^. (c) We adjust the time interval to compute the time derivative of protein concentration and compute the slope in the linear fit of Δ*p*/Δ*t vs*. *p*. The normalized slope is linearly fitted as a function of the inverse of the time interval. The fraction of intrinsic noise in the total noise can be calculated from the intercept of the linear fit. We also infer the fraction of measurement noise in the total noise from the slope of the linear fit. (d) We summarize the calculated fractions of different noise for the two data sets. *f*_in_: the fraction of intrinsic noise. *f*_ex_: the fraction of extrinsic noise. *f*_me_: the fraction of measurement noise.

In this way we find that the ratio between the measurement noise and the total noise in the two data sets are respectively 17% and 10% in terms of their standard deviations. We can further use our analytic results for constitutively expressed genes as used in these experiments to estimate the average copy numbers of proteins at cell birth and the translation burst parameter *βτ*_*m*_ (see Eqs. S28, S29 in SI C) [30]. We find that *N*_*p*_ ≈ 230 at cell birth, *βτ*_*m*_ ≈ 1.37 for Data in Fig. 4(a), and Np ≈ 210 at cell birth, *βτ*_*m*_ ≈ 2.81 for Data in Fig. 4(b).

### Summary and outlook

In this work, we start from a general framework of stochastic gene expression in exponentially growing cells. Our approach allows us to take into account the cell growth and division explicitly and study the variability in protein *concentrations*, directly relevant to experiments on proliferating cells such as bacteria, yeast or cancer cells. We derive a broadly applicable decomposition of the protein concentration noise, finding that the total noise can be expressed as the sum of the noise due to upstream factors, the Poisson noise due to the random process of production and degradation, and the noise due to random partitioning during cell division. These results are independent of the underlying details of the particular dynamics of mRNA and protein synthesis. Given a time trajectory of protein concentration, one may linearly fit the protein production rate as a function of the protein concentration. We find that the slope of the fit, normalized by the growth rate, precisely equals the fraction of intrinsic noise in the total protein concentration noise in the absence of measurement noise. We verify our theoretical framework on synthetic data of protein concentrations both for constitutively expressed genes and genes with various underlying gene expression dynamics.

Importantly, we generalize our protocol to analyze experimental data of *E. coli* gene expression and show how a generalization of the method can simultaneously reveal the fraction of *measurement noise* in addition to that of intrinsic and extrinsic noise. Our framework predicts that the slope in the linear fitting of the time derivative of protein concentration *vs*. the current protein concentration has a linear dependence on the inverse of the time-interval used in the experiments, which agrees well with the experimental results. Assuming a model of a constitutively expressed protein as used in these experiments, our approach also allows us to infer the average copy numbers of proteins at cell birth as well as the translation burst parameter, relying only on time-series data of protein concentrations in proliferating cells.

The generality of our approach and the agreement between experiments and theoretical predictions suggests that the method should be broadly applicable and will serve as a useful tool for gene expression analysis. Our protocol to extract the intrinsic and extrinsic noise relies only on the time trajectory of protein concentration of a single gene, in contrast to the dual-reporter protocol which relies on measuring protein concentrations of two identical genes. Combing our method with the dual-reporter method, one can further decompose the extrinsic noise into correlated and uncorrelated components. Theoretically, our work elucidates how various processes contribute to the gene expression noise in proliferating cells, and paves the way to further studies on the nature of the widely-observed yet poorly understood extrinsic noise in gene expression.

We thank Ido Golding and Lydia Robert for useful discussions and feedback. A.A. was supported by NSF CAREER grant 1752024 and the Harvard Dean’s Competitive Fund. A.A. and J.L. thank support from Harvard’s MRSEC (DMR-1420570).

