## Supplementary Information for "Disentangling intrinsic and extrinsic gene expression noise in growing cells"

Jie Lin and Ariel Amir
*John A. Paulson School of Engineering and Applied Sciences, Harvard University, Cambridge, MA 02138, USA*
(Dated: August 23, 2020)

### A. Derivation of the decomposition of noise

For any chemical reaction in which the number of particle $i$ changes $d_{\alpha,i}$ in reaction $\alpha$ with rate $r_\alpha$ [1], the time dependence of the covariance of the numbers of particle $i$ and $j$ is

$$\frac{d\text{cov}(x_i, x_j)}{dt} = \sum_\alpha d_{\alpha,i}(\overline{x_j r_\alpha} - \overline{x_j}\ \overline{r_\alpha}) + d_{\alpha,j}((\overline{x_i r_\alpha} - \overline{x_i}\ \overline{r_\alpha}) + d_{\alpha,i} d_{\alpha,j} \overline{r_\alpha}. \tag{S1}$$

Here $\text{cov}(x, y) = \overline{xy} - \overline{x}\ \overline{y}$. Based on the chemical reactions of gene expression we introduced in Fig. 1 in the main text,

$$N_m \xrightarrow{k_1 V} N_m + 1, \tag{S2}$$

$$N_m \xrightarrow{N_m/\tau_m} N_m - 1, \tag{S3}$$

$$N_p \xrightarrow{k_2 V} N_p + 1, \tag{S4}$$

one can find the time-dependence of the variance of protein concentration using $p = N_p/V(\tau)$:

$$\frac{d\sigma_p^2(\tau)}{d\tau} = 2\text{cov}(k_2(\tau), p) + \frac{\overline{k_2}(\tau)}{V(\tau)} - 2\mu\sigma_p^2(\tau). \tag{S5}$$

Here we use $\sigma_p^2(\tau)$ to represent the *instantaneous* variances of protein concentrations as functions of the age. The average $\overline{x}(\tau)$ is over different cell cycles conditioned on the same age, $\sigma_x^2(\tau) = \overline{x^2} - \overline{x}^2$, and $\text{cov}(x, y)(\tau) = \overline{xy} - \overline{x}\ \overline{y}$. We consider a uniform sampled data of protein concentration in time and find the instantaneous variances of protein concentrations using Eq. S5 which we sum over uniformly in time within a cell cycle (note that this protocol is not valid for data from a snapshot of a population of growing cells in which the age distribution is non-uniform [2, 3]). What is left on the left side of Eq. S5 is simply proportional to $\sigma_p^2(0) - \sigma_p^2(T)$ where $T$ is the doubling time. Because of the random partitioning of molecules, the variances of protein concentrations at cell division and at cell birth are related by $\sigma_p^2(0) = \sigma_p^2(T) + \overline{p(T)}/2V_b$ [1]. Rewriting Eq. S5, we obtain the general expression of the variance of protein concentration:

$$\sigma_p^2 = \underbrace{\frac{\text{cov}(k_2, p)}{\mu}}_{\text{Upstream noise}} + \underbrace{\left\langle \frac{k_2}{2\mu V} \right\rangle}_{\text{Poisson noise}} + \underbrace{\frac{\overline{p(T)}}{4\mu V_b T}}_{\text{Partitioning noise}}. \tag{S6}$$

Here $\text{cov}(k_2, p) = \langle k_2 p \rangle - \langle k_2 \rangle \langle p \rangle$ and $\langle \cdot\cdot \rangle$ represents average over time.

Using Eq. S1, we also obtain the time-dependence of the covariance of mRNA and protein concentrations, and the variance of mRNA concentration using $m = N_m/V(\tau)$, $p = N_p/V(\tau)$:

$$\frac{d\text{cov}(m, p)(\tau)}{d\tau} = \text{cov}(k_2, m) + \text{cov}(k_1, p) - (2\mu + \frac{1}{\tau_m})\text{cov}(m, p)(\tau), \tag{S7}$$

$$\frac{d\sigma_m^2(\tau)}{d\tau} = 2\text{cov}(k_1, m) + \frac{\overline{m}}{\tau_m V(\tau)} + \frac{\overline{k_1}}{V(\tau)} - 2(\mu + \frac{1}{\tau_m})\sigma_m^2(\tau). \tag{S8}$$

Here we use $\sigma_m^2(\tau)$ to represent the *instantaneous* variance of mRNA concentration as a function of the age. The average $\overline{x}(\tau)$ is over different cell cycles conditioned on the same age, $\sigma_x^2(\tau) = \overline{x^2} - \overline{x}^2$, and $\text{cov}(x, y)(\tau) = \overline{xy} - \overline{x}\ \overline{y}$.

Using a similar argument for mRNAs, we sum over Eq. S8 uniformly in time with a small time interval $\Delta t$ within a cell cycle. Using the boundary condition of mRNA concentration, we obtain

$$\sigma_m^2 = \underbrace{\frac{\text{cov}(k_1, m)}{\mu + \frac{1}{\tau_m}}}_{\text{Upstream noise}} + \underbrace{\left\langle \frac{\frac{m(t)}{\tau_m} + k_1(t)}{2(\mu + \frac{1}{\tau_m})V(t)} \right\rangle}_{\text{Poisson noise}} + \underbrace{\frac{\langle m(T) \rangle}{4(\mu + \frac{1}{\tau_m}))V_b T}}_{\text{Partitioning noise}}. \tag{S9}$$

Similar logic leads to the following equation for proteins with a finite lifetime $\tau_p$,

$$\sigma_p^2 = \underbrace{\frac{\mathrm{cov}(k_2, m)}{\mu + \frac{1}{\tau_p}}}_{\text{Upstream noise}} + \underbrace{\left\langle \frac{\frac{p(t)}{\tau_p} + k_2(t)}{2(\mu + \frac{1}{\tau_p})V(t)} \right\rangle}_{\text{Poisson noise}} + \underbrace{\frac{\langle p(T) \rangle}{4(\mu + \frac{1}{\tau_p}))V_b T}}_{\text{Partitioning noise}}. \tag{S10}$$

Our protocol to extract the fraction of intrinsic noise is equally valid for a degradable proteins given its lifetime $\tau_p$ is known. In this case, one should replace $\mu$ by $\mu + 1/\tau_p$ in Eqs. 4, 6 in the main text. If the cell does not grow, $\mu = 0$ and the intrinsic noise only includes the Poisson noise.

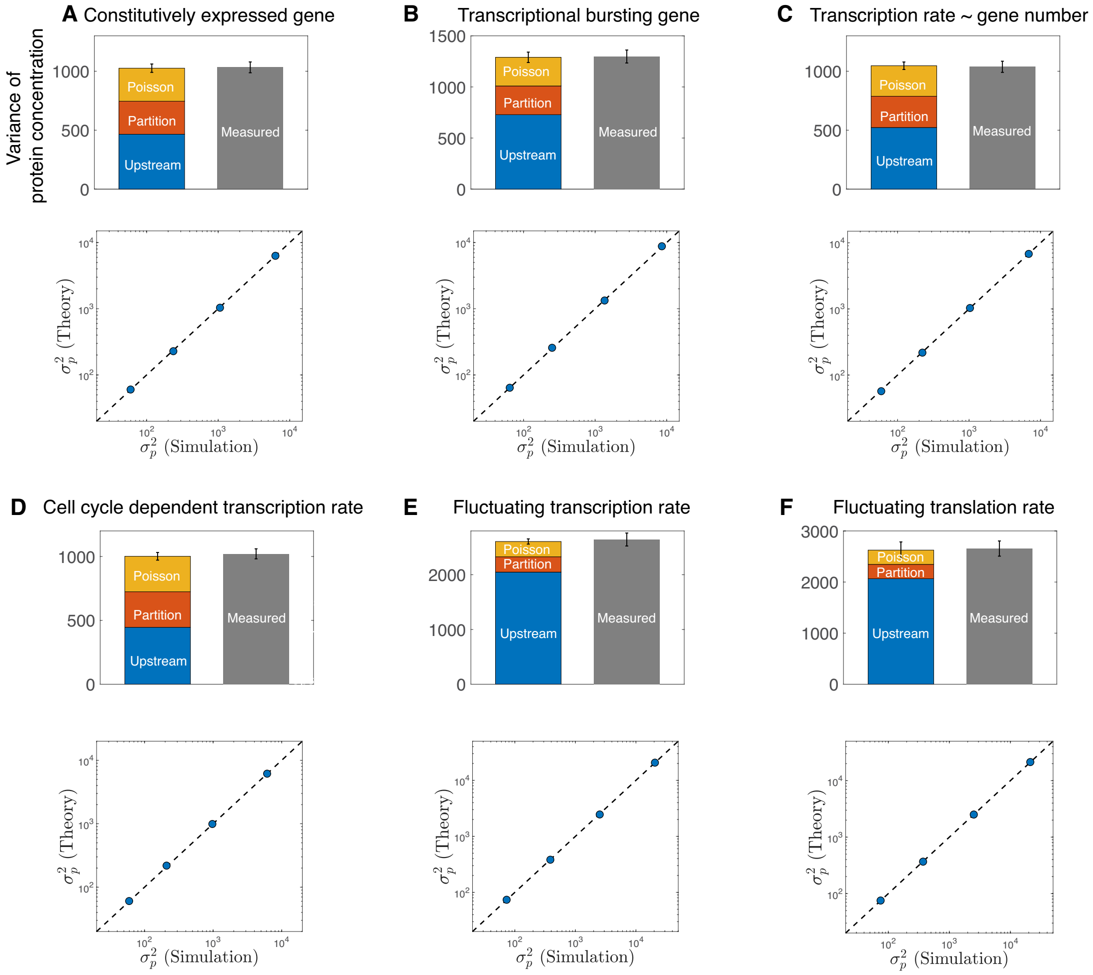


**Supplementary Figure S1 Decomposition of the noise of protein concentration.**
(**A**) Simulation of a constitutively expressed gene. (Upper) Total measured noise of protein concentration and the three sources of noise. (Bottom) The translation rate per mRNA $\beta$ is varied and the predicted variance of protein concentration (Eq. 1 in the main text) is compared with the measured value. The same analysis applies to the following panels. (**B**) Simulation of a transcriptional bursting gene. (**C**) Simulation of a scenario where the transcription rate is proportional to the gene copy number. (**D**) Simulation of a gene with transcription rate modulated throughout the cell cycle. (**E**) Simulation of a gene with a fluctuating transcription rate such that $k_1(t) = \langle k_1 \rangle + \xi_1(t)$ where $\xi_1(t)$ is the noise term. (**F**) Simulation of a gene with a fluctuating translation rate per mRNA such that $\beta(t) = \langle \beta \rangle + \xi_2(t)$ where $\xi_2(t)$ is the noise term. In all panels, $T = 60$, $\tau_m = 10$, and $k_1 = 10$ if not specified. In all upper panels, $\beta = 0.1$ and in all bottom panels, $\beta$ is varied so that $\log_{10} \beta = -2, -1.5, -1, -0.5$. Other simulation details are explained in the text. The errorbars are computed as the standard deviation over 5 independent simulations.

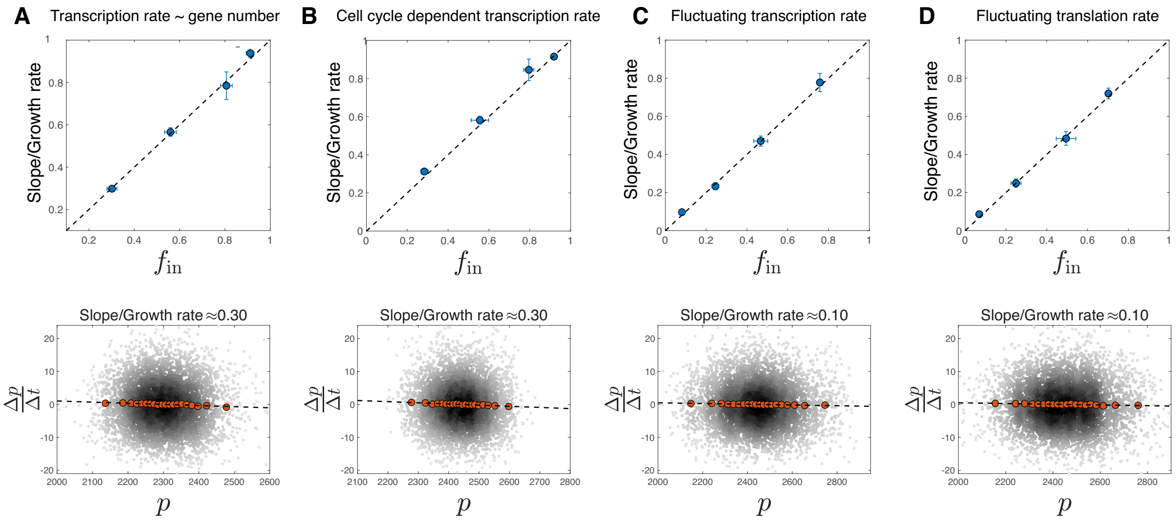


**Supplementary Figure S2 Extraction of the fraction of intrinsic noise based on synthetic data.** (**A**) Simulation of a scenario where the transcription rate is proportional to the gene copy number. (**B**) Simulation of a gene with transcription rate modulated throughout the cell cycle. (**C**) Simulation of a gene with a fluctuating transcription rate $k_1(t)$. (**D**) Simulation of a gene with a fluctuating translation rate per mRNA $\beta(t)$. In all upper panels, $\beta$ is varied so that $\log_{10}\beta = -2, -1.5, -1, -0.5$. In all bottom panels, $\log_{10}\beta = -0.5$. We compute the time-derivative of protein concentration with a time interval $\Delta t = 0.5$. Other simulation details are explained in Methods. The errorbars are computed as the standard deviation of 5 independent simulations.

### B. Numerical confirmation of the noise decomposition and extraction of intrinsic noise

We test the decomposition of noise for protein concentration, Eq. 1 in the main text by simulating genes with various underlying dynamics. In all panels of Fig. S1, $T = 60$, $\tau_m = 10$, and $k_1 = 10$ if not specified. In Fig. S1A, we simulate a constitutively expressed gene with constant transcription rate $k_1$ and constant translation rate per mRNA $\beta$. In Fig. S1B, we simulate a gene with transcriptional bursting such that transcription only occurs in the "on" state. $k_{\text{on}} = 10$, $k_{\text{off}} = 10$, $k_1 = 20$. In Fig. S1C, we simulate a gene with a transcription rate proportional to the gene copy number which doubles in the middle of the cell cycle. The transcription rate changes from $k_1$ to $2k_1$ in the middle of the cell cycle and $k_1$ is chosen such that the average mRNA number at cell birth is the same as the constitutively expressed gene. In Fig. S1D, we simulate a gene with a transcription rate that depends on the cell cycle due to a finite period of DNA replication (section E). In Fig. S1E, we simulate a gene with a fluctuating transcription rate such that $k_1(t) = \langle k_1 \rangle + \xi_1(t)$ and the autocorrelation function of the noise decays exponentially in time, $\langle \xi_1(t)\xi_1(t') \rangle = A_1 \exp(-|t-t'|/\tau_1)$. We take $A_1 = 0.01k_1^2$ and $\tau_1 = T/2$ in the simulation where $T$ is the cell cycle duration. In Fig. S1F, we simulate a gene with a fluctuating translation rate per mRNA such that $\beta = \langle \beta \rangle + \xi_2(t)$ where $\langle \xi_2(t)\xi_2(t') \rangle = A_2 \exp(-|t-t'|/\tau_2)$. We take $A_2 = 0.01\langle \beta \rangle^2$ and $\tau_2 = T/2$ in the simulation. $\log_{10}\langle \beta \rangle = -2, -1.5, -1, -0.5$ in the upper panel of Fig. S1F. $\log_{10}\langle \beta \rangle = -0.5$ in the bottom panel of Fig. S1F. The same simulations discussed above are used to confirm the validity of extraction of intrinsic noise, Eq. 4 in the main text, shown in Fig. S2.

### C. Mathematical derivations for a constitutively expressed gene

Using the fact that $k_1$ is constant for a constitutively expressed gene, Eqs. 4, 13, 14 in the main text are simplified to

$$\frac{d\sigma_m^2}{d\tau} = \frac{\langle m \rangle}{\tau_m} + \frac{k_1}{V(\tau)} - 2(\mu + \frac{1}{\tau_m})\sigma_m^2, \tag{S11}$$

$$\frac{d\text{cov}(m,p)}{d\tau} = \beta\sigma_m^2 - (2\mu + \frac{1}{\tau_m})\text{cov}(m,p), \tag{S12}$$

$$\frac{d\sigma_p^2}{d\tau} = 2\beta\text{cov}(m,p) + \frac{\beta\langle m \rangle}{V(\tau)} - 2\mu\sigma_p^2, \tag{S13}$$

where $\langle m\rangle = k_1/(\mu + 1/\tau_m)$. We solve Eq. (S11) relying on the exponential growth of cell volume $V(\tau) = V_b e^{\mu\tau}$ and find

$$\sigma_m^2(\tau) = \sigma_m^2(0)e^{-2(\mu+\frac{1}{\tau_m})\tau} + \frac{k_1 + \frac{\langle m\rangle}{\tau_m}}{V_b}\frac{1}{\mu + \frac{2}{\tau_m}}(e^{-\mu\tau} - e^{-2(\mu+\frac{1}{\tau_m})\tau}). \tag{S14}$$

We now take $\tau = \log(2)/\mu$ and use the boundary condition $\sigma_m^2(0) - \sigma_m^2(T) = \langle m\rangle/(2V_b)$ to obtain

$$\sigma_m^2(T) = \sigma_m^2(0)2^{-2(1+\frac{1}{\mu\tau_m})} + \frac{k_1 + \frac{\langle m\rangle}{\tau_m}}{V_b}\frac{1}{\mu + \frac{2}{\tau_m}}\Big(\frac{1}{2} - \big(\frac{1}{2}\big)^{2+\frac{2}{\mu\tau_m}}\Big) = \sigma_m^2(0) - \frac{\langle m\rangle}{2V_b}, \tag{S15}$$

from which we obtain

$$\sigma_m^2(0) = \frac{\frac{k_1+\frac{\langle m\rangle}{\tau_m}}{V_b}\frac{1}{\mu+\frac{2}{\tau_m}}\Big(\frac{1}{2} - \big(\frac{1}{2}\big)^{2+\frac{2}{\mu\tau_m}}\Big) + \frac{\langle m\rangle}{2V_b}}{1 - 2^{-2(1+\frac{1}{\mu\tau_m})}}. \tag{S16}$$

Using the expression of $\langle m\rangle$, it is straightforward to verify that $\frac{k_1+\frac{\langle m\rangle}{\tau_m}}{\mu+\frac{2}{\tau_m}} = \langle m\rangle$, which leads to

$$\sigma_m^2(\tau) = \frac{\langle m\rangle}{V_b}e^{-\mu\tau}. \tag{S17}$$

Given the time dependence of $\sigma_m^2(\tau)$ we can further solve for $\mathrm{cov}(m,p)$ using Eq. S12 and find the general solution as

$$\mathrm{cov}(m,p) = \mathrm{cov}(m,p)(\tau=0)e^{-(2\mu+\frac{1}{\tau_m})\tau} + \frac{\beta\langle m\rangle}{V_b}\frac{1}{\mu+\frac{1}{\tau_m}}\Big(e^{-\mu\tau} - e^{-(2\mu+\frac{1}{\tau_m})\tau}\Big). \tag{S18}$$

Using the boundary condition $\mathrm{cov}(m,p)(\tau=0) = \mathrm{cov}(m,p)(\tau=T)$, we find that

$$\mathrm{cov}(m,p)(\tau=0) = \frac{\beta\langle m\rangle}{V_b}\frac{1}{\mu+\frac{1}{\tau_m}}\frac{\frac{1}{2} - (\frac{1}{2})^{2+\frac{1}{\mu\tau_m}}}{1 - (\frac{1}{2})^{2+\frac{1}{\mu\tau_m}}}. \tag{S19}$$

Therefore,

$$\begin{aligned}\mathrm{cov}(m,p) &= \frac{\beta\langle m\rangle}{V_b}\frac{1}{\mu+\frac{1}{\tau_m}}\frac{\frac{1}{2} - (\frac{1}{2})^{2+\frac{1}{\mu\tau_m}}}{1 - (\frac{1}{2})^{2+\frac{1}{\mu\tau_m}}}e^{-(2\mu+\frac{1}{\tau_m})\tau} + \frac{\beta\langle m\rangle}{V_b}\frac{1}{\mu+\frac{1}{\tau_m}}\Big(e^{-\mu\tau} - e^{-(2\mu+\frac{1}{\tau_m})\tau}\Big) \\ &= -\frac{\beta\langle m\rangle}{V_b}\frac{1}{\mu+\frac{1}{\tau_m}}\frac{2^{1+\frac{1}{\mu\tau_m}}}{2^{2+\frac{1}{\mu\tau_m}} - 1}e^{-(2\mu+\frac{1}{\tau_m})\tau} + \frac{\beta\langle m\rangle}{V_b}\frac{1}{\mu+\frac{1}{\tau_m}}e^{-\mu\tau} \\ &= Ae^{-(2\mu+\frac{1}{\tau_m})\tau} + Be^{-\mu\tau},\end{aligned} \tag{S20}$$

with $A = -\frac{\beta\langle m\rangle}{V_b}\frac{1}{\mu+\frac{1}{\tau_m}}\frac{2^{1+\frac{1}{\mu\tau_m}}}{2^{2+\frac{1}{\mu\tau_m}}-1}$ and $B = \frac{\beta\langle m\rangle}{V_b}\frac{1}{\mu+\frac{1}{\tau_m}}$. Given the time dependence of $\mathrm{cov}(m,p)$ we can rewrite Eq. (S13) as

$$\frac{d\sigma_p^2}{dt} = 2\beta Ae^{-(2\mu+\frac{1}{\tau_m})\tau} + 2\beta Be^{-\mu\tau} + \frac{\beta\langle m\rangle}{V_b}e^{-\mu\tau} - 2\mu\sigma_p^2. \tag{S21}$$

Its general solution is

$$\sigma_p^2(\tau) = \sigma_p^2(0)e^{-2\mu\tau} + 2\beta A\tau_m(e^{-2\mu\tau} - e^{-(2\mu+\frac{1}{\tau_m})\tau}) + \frac{2\beta B + \frac{\beta\langle m\rangle}{V_b}}{\mu}(e^{-\mu\tau} - e^{-2\mu\tau}). \tag{S22}$$

We now take $\tau = \log(2)/\mu$ and use the boundary condition $\sigma_p^2(0) - \sigma_p^2(T) = \langle p\rangle/(2V_b)$ so that

$$\sigma_p^2(T) = \frac{\sigma_p^2(0)}{4} + 2\beta A\tau_m\Big(\frac{1}{4} - \big(\frac{1}{2}\big)^{2+\frac{1}{\mu\tau_m}}\Big) + \frac{2\beta B + \frac{\beta\langle m\rangle}{V_b}}{4\mu} = \sigma_p^2(0) - \frac{\langle p\rangle}{2V_b}. \tag{S23}$$

We find that

$$\sigma_p^2(0) = \frac{2\langle p\rangle}{3V_b} + \frac{2\beta A\tau_m}{3}\Big(1-\big(\frac{1}{2}\big)^{\frac{1}{\mu\tau_m}}\Big) + \frac{2\beta B + \frac{\beta\langle m\rangle}{V_b}}{3\mu} = \frac{\langle p\rangle}{V_b} + \frac{2\beta A\tau_m}{3}\Big(1-\big(\frac{1}{2}\big)^{\frac{1}{\mu\tau_m}}\Big) + \frac{2\beta B}{3\mu} \tag{S24}$$

where we have used $\langle p\rangle = \beta\langle m\rangle/\mu$.

We now compute the upstream noise, the Poisson noise and the partitioning noise for a constitutively expressed gene using Eq. S6. In this case, the Poisson noise is equal to the partitioning noise and

$$\sigma^2_{p,\text{poisson}} = \sigma^2_{p,\text{partitioning}} = \frac{\langle p\rangle}{4\ln(2)V_b}. \tag{S25}$$

In the limit $\mu\tau_m \ll 1$, we find

$$\text{cov}(m,p) = \frac{\beta\langle m\rangle\tau_m}{V_b}e^{-\mu\tau}. \tag{S26}$$

Therefore, the upstream noise becomes

$$\sigma^2_{p,\text{upstream}} = \frac{\beta\int_0^T \text{cov}(m,p)dt}{T\mu} = \frac{\langle p\rangle}{2\ln(2)}\frac{\beta\tau_m}{V_b}. \tag{S27}$$

We can also rewrite Eqs. S25, S27 in terms of $\text{CV}^2$ (variance/mean$^2$) as

$$\text{CV}^2_{\text{intrinsic}} = \frac{1}{2\ln(2)\langle N_{p,b}\rangle}, \tag{S28}$$

$$\text{CV}^2_{\text{extrinsic}} = \frac{\beta\tau_m}{2\ln(2)\langle N_{p,b}\rangle}, \tag{S29}$$

here $\langle N_{p,b}\rangle$ is the average protein number at cell birth. The above calculation can be easily generalized to a constitutively expressed protein with a finite lifetime $\tau_p$. Eqs. S12, S13 are modified as

$$\frac{d\text{cov}(m,p)}{d\tau} = \beta\sigma_m^2 - (2\mu + \frac{1}{\tau_m} + \frac{1}{\tau_p})\text{cov}(m,p), \tag{S30}$$

$$\frac{d\sigma_p^2}{d\tau} = 2\beta\text{cov}(m,p) + \frac{\beta\langle m\rangle}{V(\tau)} - 2(\mu + \frac{1}{\tau_p})\sigma_p^2. \tag{S31}$$

Therefore the solution of Eq. (S12) is valid for Eq. (S30) as well after replacing $1/\tau_m$ by $1/\tau_m + 1/\tau_p$. In the limit $\mu\tau_m \ll 1$, we find

$$\text{cov}(m,p) = \frac{\beta\langle m\rangle\tau_m\tau_p}{V_b(\tau_m+\tau_p)}e^{-\mu\tau}. \tag{S32}$$

Using Eq. (16) in the maintext, the upstream noise becomes

$$\sigma^2_{p,\text{upstream}} = \frac{\beta\int_0^T \text{cov}(m,p)dt}{T(\mu+\frac{1}{\tau_p})} = \frac{\langle p\rangle}{2\ln(2)(1+\frac{1}{\mu\tau_p})}\frac{\beta\tau_m\tau_p}{V_b(\tau_m+\tau_p)}. \tag{S33}$$

Similarly, it is straightforward to find that in this case

$$\sigma^2_{p,\text{poisson}} = \frac{\langle p\rangle(1+\frac{2}{\mu\tau_p})}{4\ln(2)(1+\frac{1}{\mu\tau_p})V_b}, \tag{S34}$$

$$\sigma^2_{p,\text{partitioning}} = \frac{\langle p\rangle}{4\ln(2)(1+\frac{1}{\mu\tau_p})V_b}. \tag{S35}$$

#### D. Noise strength of a constitutively expressed gene

To quantify the noise strength of protein concentration, one can either use the $\text{CV}^2$ (variance/mean$^2$) or the Fano factor (variance/mean). While the $\text{CV}^2$ is dimensionless, the Fano factor is dimensional. To make the Fano factor of protein concentration dimensionless, one needs to multiply the Fano factor of protein concentration by an arbitrary volume scale $\tilde{V}$. A common way is to use the average cell volume $\overline{V}$ [4]. For uniform-time sampling, the average cell volume is simply $\overline{V} = \int_0^T V_b e^{\mu t} dt/T = V_b/\ln 2$. Assuming a non-degradable protein and $\mu\tau_m \ll 1$, we obtain

$$\text{CV}^2 = \frac{1+\beta\tau_m}{2\ln(2)\langle p\rangle V_b}, \tag{S36}$$

$$\text{Fano} = \frac{1+\beta\tau_m}{2\ln 2 V_b}\overline{V} = \frac{1+\beta\tau_m}{2(\ln 2)^2} \approx 1.04(1+\beta\tau_m). \tag{S37}$$

Experimentally, it is also common to sample the data by taking a snapshot of a population of cells. In this protocol, the distribution of age is non-uniform and decays exponentially, $P(\tau) = 2\mu e^{-\mu\tau}$. The average cell volume is $\overline{V} = \int_{V_b}^{2V_b} V \frac{2}{V^2} dV = 2\ln 2 V_b$. Using Eq. S22, we find that

$$\text{CV}^2 = \frac{13}{18}\frac{\frac{27}{26}+\beta\tau_m}{\langle p\rangle V_b} \approx 0.72\frac{1.04+\kappa_2\tau_1}{\langle p\rangle V_b}, \tag{S38}$$

$$\text{Fano} = \frac{13}{18}\frac{\frac{27}{26}+\beta\tau_m}{V_b}\overline{V} \approx 1.04+\beta\tau_m. \tag{S39}$$

Interestingly, the Fano factors derived here are very close to the Fano factor $1+\beta\tau_m$ one would obtain from the constant cell volume model [5].

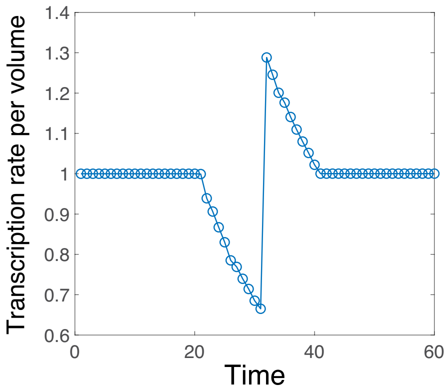


**Supplementary Figure S3 Numerical simulation of a cell cycle with a finite DNA replication period.** The time dependence of the transcription rate per cell volume $k_1$.

#### E. Cell cycle with a finite DNA replication period

We relax the assumption of instantaneous DNA replication in the main text to take into account the effects of a finite DNA replication period on the transcription rate per cell volume $k_1$. We assume a doubling time of 60 mins. The gene is constitutively expressed and replicated in the middle of the cell cycle at $\tau = 30$ mins. DNA replication starts from $\tau = 20$ mins with a duration 20 mins. Because of the competition between genes for the limiting resource such as RNA polymerase [6, 7], the transcription rate of the gene under consideration decreases during DNA replication with a jump right after the gene is duplicated (Fig. S3). Our predictions regarding the decomposition of the noise of protein concentration and its magnitude (Eq. 5 in the main text), and the extraction of intrinsic noise (Eq. 9 in the main text) are nicely confirmed in this case as well (Fig. 2D and Fig. 3D in the main text).

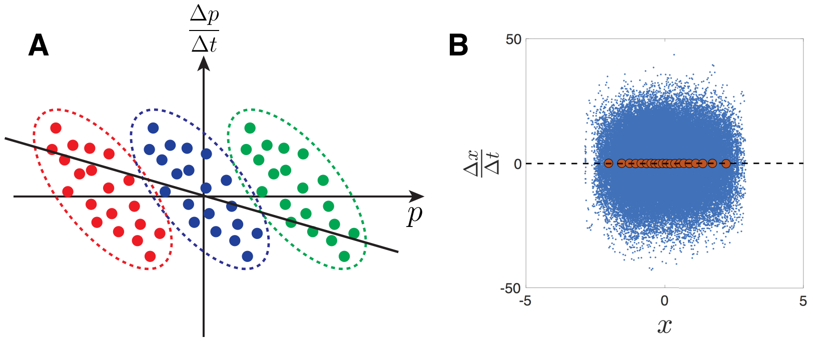


**Supplementary Figure S4 Simplified model based on the Langevin equation.** (**A**) Schematic illustration of the lowering of the slope due to extrinsic noise; In essence, this is due to Simpson's paradox [8]. (**B**) Numerical test of the simplified model. Here, $\mu = 1$, $\tau_\eta = 0.01$, $A_\eta = 100$, $D_\xi = 0$ and the time interval of the simulation $\Delta t = 0.001$. The red circles are binned data and the dashed line has a zero slope.

#### F. Simplified model based on the Langevin equation

To better understand the effects of extrinsic noise, we consider a simplified version of the model in the main text by taking a small time interval such that $k_2 V \delta t \gg 1$ but also small enough that the change of $k_2$ is negligible. On such a time scale, the chemical reaction Eq. 3 introduced in the main text can be approximated by a discrete Langevin equation [9]:

$$\delta p = (k_2 - \mu p)\delta t + \sqrt{\frac{k_2 \delta t}{V}} W. \tag{S40}$$

Here the noise stemming from the random production becomes white noise where $W$ is a Gaussian random variable with variance 1. For simplicity, we have neglected the partitioning noise. Since the time interval $\delta t$ can be arbitrarily small in the large volume limit, the upstream noise in the translation rate $k_2$ should be considered as a continuous random variable (i.e., it has a finite correlation time). Finally, we obtain the following continuous Langevin equation as an approximation for the dynamics of protein concentration:

$$\dot{x} = -\mu x + \eta + \xi, \tag{S41}$$

where $x$ is the deviation of the protein concentration from its average. $\xi$ is a white noise term such that $\langle \xi(t)\xi(t') \rangle = 2D_\xi \delta(t - t')$. $\eta$ corresponds to the upstream noise in the translation rate which is a continuous random variable. For concreteness, we assume the autocorrelation function of $\eta$ decays exponentially so that $\langle \eta(t)\eta(t') \rangle = A_\eta e^{-|t-t'|/\tau_\eta}$.

Before we move on to the theoretical analysis of the above equation, we first present an intuitive argument why the extrinsic noise reduces the slope in the linear fitting of $\Delta x/\Delta t$ vs. $x$. Consider the limit $\tau_\eta \gg 1/\mu$ so that $\eta$ can be approximated as a constant for many generations of cell cycle. For a constant $\eta$ the slope is simply the growth rate $\mu$. Combining multiple sets of data with different $\eta$'s, it is evident that the slope becomes smaller than $\mu$ (Fig. S4A). Note that this effect is essentially due to "Simpson's paradox", where correlations can dramatically change when pooling together different sub-populations [8, 10]. While this argument is based on the assumption of a very slow time dependence of $\eta$, the conclusion is generally valid as we show in the following.

From Eq. S41, the solution of $x$ can be formally written as

$$x = \int_0^t e^{-\mu(t-t')}[\eta(t') + \xi(t')]dt', \tag{S42}$$

where the term associated with the initial condition is neglected since we are interested in the steady state. It is straightforward to find the variance of $x$ as

$$\sigma_x^2 = \frac{A_\eta \tau_\eta}{\mu} \frac{1}{1 + \mu\tau_\eta} + \frac{D_\xi}{\mu}, \tag{S43}$$

where the first term can be considered as the upstream noise in the main text and the second term can be considered as the intrinsic noise. We now calculate the covariance between $\frac{\Delta x}{\Delta t}$ and $x$:

$$\mathrm{cov}(\frac{\Delta x}{\Delta t}, x) = -\mu\sigma_x^2 + \mathrm{cov}(\eta, x). \tag{S44}$$

Therefore, the slope (S) in the linear regression of $\frac{\Delta x}{\Delta t}$ *vs.* $x$ becomes

$$S \equiv -\frac{\mathrm{cov}(\frac{\Delta x}{\Delta t}, x)}{\sigma_x^2} = \mu - \frac{\mathrm{cov}(\eta, x)}{\sigma_x^2}. \tag{S45}$$

We now calculate $\mathrm{cov}(\eta, x)$

$$\mathrm{cov}(\eta, x) = \int_0^t e^{-\mu(t-t')} \langle \eta(t)\eta(t') \rangle dt' = \frac{A_\eta \tau_\eta}{1 + \mu \tau_\eta}$$

Combining with Eq. (S43, S45), we find that the relative slope indeed tells us the relative fraction of intrinsic noise in the total noise

$$\frac{S}{\mu} = \frac{D_\xi/\mu}{\sigma_x^2}. \tag{S46}$$

Note that when $D_\eta > 0$ and $D_\xi = 0$ the noise of $x$ is all coming from $\eta$ and $S = 0$, while the variance of $x$ is still finite. The above calculation is valid even in the limit $\tau_\eta \ll 1/\mu$, which is beyond the simple argument assuming $\tau_\eta \gg 1/\mu$. We numerically test this situation and find a zero slope in the linear fitting of $\Delta x/\Delta t$ *v.s.* $x$ (Fig. S4B).

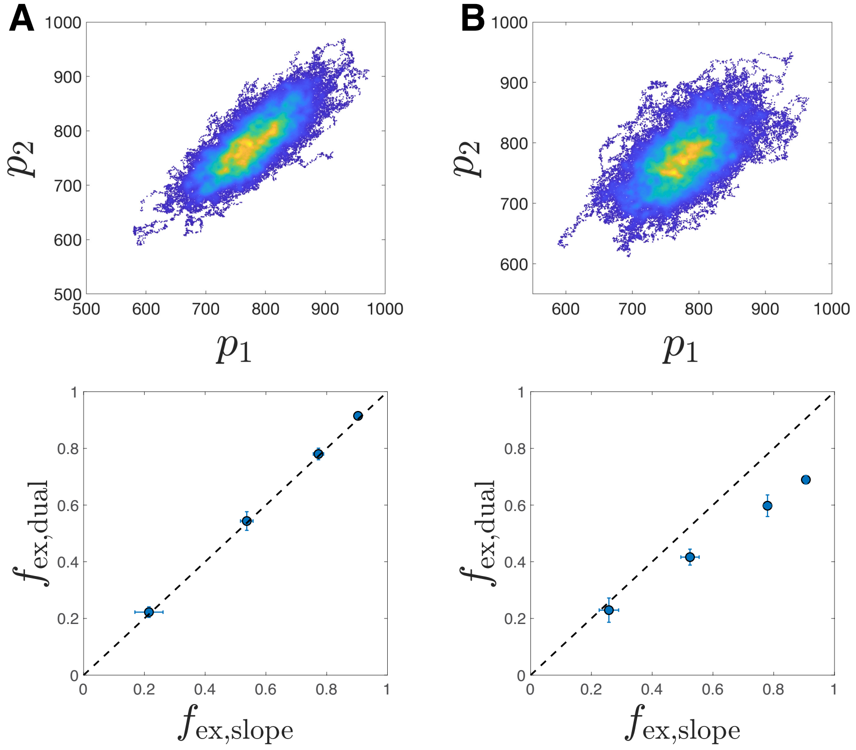


**Supplementary Figure S5 Numerical test of dual-reporter setup.**
(**A**) We simulate two identical genes that share the same fluctuating translation rate $k_2(t)$ and set the translation rate per mRNA as $\beta = \langle \beta \rangle + \xi_2(t)$ where $\xi_2(t)$ is the noise term. We assume $\langle \xi_2(t)\xi_2(t') \rangle = A_2 \exp(-|t-t'|/\tau_2)$ with $A_2 = 0.01\beta^2$ and $\tau_2 = T/2$. $T = 60$, $k_1 = 10$ and $\tau_m = 10$. (Upper) we show the original data of the two identical genes with $\langle \beta \rangle = 0.1$. (Bottom) We compare the fractions of extrinsic noise inferred from the dual-reporter setup and from the slope in the linear fitting of $\Delta p/\Delta t$ *vs.* $p$. $\beta$ is varied so that $\log_{10}\langle \beta \rangle = -2, -1.5, -1, -0.5$. (**B**) We simulate two identical genes that share the same fluctuating transcription rate $k_1(t)$ and set the transcription rate as $k_1(t) = \langle k_1 \rangle + \xi_1(t)$ where $\xi_1(t)$ is the noise term. We assume $\langle \xi_1(t)\xi_1(t') \rangle = A_1 \exp(-|t-t'|/\tau_1)$ with $A_1 = 0.01k_1^2$ and $\tau_1 = T/2$. $T = 60$, $\langle k_1 \rangle = 10$ and $\tau_m = 10$. (Upper) we show the original data of the two identical genes with $\beta = 0.1$. (Bottom) $\beta$ is varied so that $\log_{10}\beta = -2, -1.5, -1, -0.5$.

#### G. Comparison with the dual-reporter setup

As we discuss in the main text, the extrinsic noise inferred from the dual-reporter setup is in general smaller than the one inferred from the time trajectory of protein concentration based on our protocol. We first consider two identical genes that share the same fluctuating translation rates $k_2(t)$ and compute the uncorrelated noise as $\sigma^2_{p,\text{uncorrelated}} = \langle (p_1(t) - p_2(t))^2 \rangle /2$ (which is referred as the intrinsic noise in the dual-reporter setup). We then compute the fraction of correlated noise as $f_{\text{ex, dual}} = (\sigma_p^2 - \sigma^2_{p,\text{uncorrelated}})/\sigma_p^2$ (which is referred as the extrinsic noise in the dual-reporter setup). We compare it with the one inferred from the slope in the linear fitting of $\Delta p/\Delta t$ *vs.* $p$, $f_{\text{ex, slope}}$ and get very nice agreement (Fig. S5A).

We also consider two identical genes that share the same fluctuating transcription rates $k_1(t)$. In this scenario, the translation rates $k_2(t)$ of the two genes are correlated but not identical due to the randomness of mRNA production and degradation. Therefore, $f_{\text{ex, dual}} < f_{\text{ex, slope}}$ as we confirm numerically (Fig. S5B).

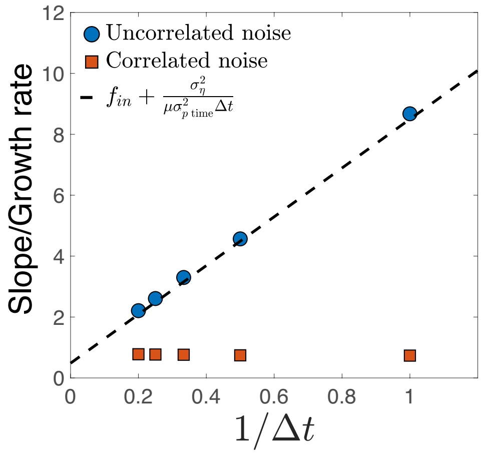


**Supplementary Figure S6 Numerical test of synthetic data with artificial measurement noise.** We simulate a constitutively expressed gene and add Gaussian noise to every recorded protein concentration (circles). $T = 60$, $k_1 = 10$, $\beta = 0.1$, $\tau_m = 10$. The time interval $\Delta t$ is varied from 1 to 5 and for each time interval we compute the slope from the linear fitting of $\Delta p/\Delta t$ *vs.* $p$. The dashed line is the theoretical prediction (Eq. 11 in the main text). $f_{\text{in}} \approx 0.50$, $\sigma^2_\eta = 100$, and $\sigma^2_{p,\text{time}} \approx 1.14 \times 10^3$. We repeat the analysis for correlated measurement noise (squares). The autocorrelation function of the measurement noise decays exponentially with a decay time $\tau_\eta = T/2$ with other parameters kept the same as uncorrelated noise.

#### H. Numerical simulation with artificial measurement noise

We simulate a constitutively expressed gene and add measurement noise to the protein concentration. We first consider the case of uncorrelated measurement noise. We vary the time interval $\Delta t$ to compute $\Delta p/\Delta t$ and compare the measured slopes in the linear fit of $\Delta p/\Delta t$ *vs.* $p$ with the theoretical prediction, Eq. 11 in the main text. The prediction is nicely confirmed (Fig. S6A). We also consider the case of correlated noise and assume the autocorrelation function of the measurement noise decays exponentially in time with a decay time $\tau_\eta$. In this case, the simulation results do not agree with Eq. (11) in the main text. The agreement of Eq. 11 in the main text and the experimental data therefore supports our assumption of uncorrelated measurement noise.

#### I. Effects of finite maturation time of fluorescent protein

We consider the effects of a finite maturation time of fluorescent protein to our theoretical predictions. We will consider two models for the maturation process: a Poisson process, following Ref. [11], and later on, a model in which the maturation process takes a constant (fixed) time. For the former, we modify Eqs. 1-3 in the main text and define

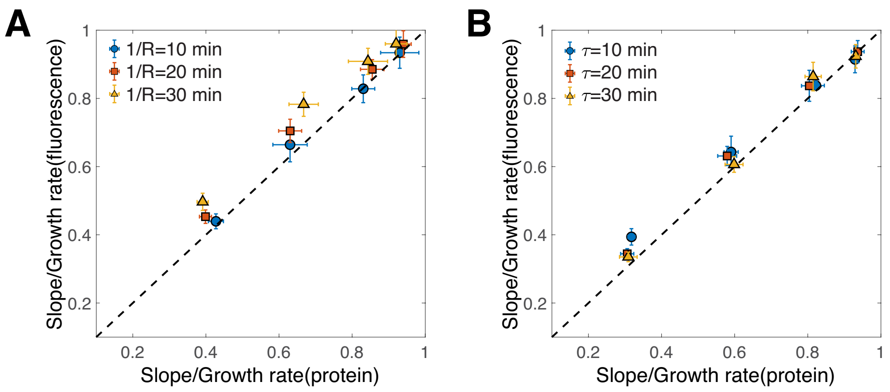


**Supplementary Figure S7 Numerical simulation of a fluorescent protein with a finite maturation time.** (**A**) We simulate a constitutively expressed gene with $T = 30$ min, $\tau_m = 10$ min, $k_1 = 10$ min$^{-1}$ and adjust $\beta$ so that $\log_{10}\beta = -2, -1.5, -1, -0.5$ min$^{-1}$. We model the maturation process as a Poisson process and compare the inferred fractions of intrinsic noise Eq. 9 in the main text and Eq. S51 here for three different maturation rates. The dashed line is the $y = x$ line. (**B**) The same analysis for a maturation process with a fixed maturation time $\tau$.

$N_u$ as the number of immature fluorescent proteins and $N_f$ as the number of maturated fluorescent proteins:

$$N_m \xrightarrow{k_1 V} N_m + 1, \tag{S47}$$

$$N_m \xrightarrow{N_m/\tau_m} N_m - 1, \tag{S48}$$

$$N_u \xrightarrow{k_2 V} N_u + 1, \tag{S49}$$

$$N_u, N_f \xrightarrow{R N_u} N_u + 1, N_f - 1. \tag{S50}$$

Here $R$ is the maturation rate. In experiments, what one can actually measure is the concentration of maturated fluorescent proteins, $f$. We compute the inferred fraction of intrinsic noise from the maturated fluorescent proteins as

$$\frac{S_f}{\mu} \equiv -\frac{\mathrm{cov}\left(\frac{\Delta f(t)}{\Delta t}, f(t)\right)}{\mu\sigma^2_{f,\mathrm{time}}}, \tag{S51}$$

and compare it to the result of Eq. 9 in the main text using the concentration of total protein $p$. We compare the two slopes ($S_f$ *vs.* $S$) using synthetic data where both can be accessed. As expected, when the maturation rate is large (corresponding to a maturation time short compared with the cell cycle duration), they are approximately equal and as the maturation rate decreases, the fraction of intrinsic noise inferred from the maturated fluorescent protein deviates from its true value, and is larger than it (Fig. S7). Typical maturation times range from several minutes to several tens of minutes [11]. The results of Fig. S7 therefore suggest that our protocol should provide accurate results for the majority of experimental scenarios.

We also consider an alternative model of maturation process, assuming that the fluorescent proteins maturate after a fixed amount of time $\tau$. We find that in this case, the inferred fractions of intrinsic noise obtained using the maturated fluorescent protein levels match closely the true values even for large maturation times. Taken together with the results for the Poisson model of protein maturation, we conclude that maturation times should not significantly affect our analysis.

---

[8] E. H. Simpson, Journal of the Royal Statistical Society: Series B (Methodological) **13**, 238 (1951).
[9] D. T. Gillespie, The Journal of Chemical Physics **113**, 297 (2000).
[10] J. Lin, M. Manhart, and A. Amir, Genetics (2020), 10.1534/genetics.120.303149.
[11] E. Balleza, J. M. Kim, and P. Cluzel, Nature Methods **15**, 47 (2018).
